# The effect of left fronto-parietal resections on hand selection: a lesion-tractography study

**DOI:** 10.1101/2019.12.11.872754

**Authors:** Henrietta Howells, Guglielmo Puglisi, Antonella Leonetti, Luca Vigano, Luca Fornia, Luciano Simone, Stephanie J. Forkel, Marco Rossi, Marco Riva, Gabriella Cerri, Lorenzo Bello

## Abstract

Strong right-hand preference on the population level is a uniquely human feature, although the neural basis for this is still not clearly defined. Recent behavioural and neuroimaging literature suggests that hand preference may be related to the orchestrated function and size of fronto-parietal white matter tracts bilaterally. Lesions to these tracts induced during tumour resection may provide an opportunity to test this hypothesis. In the present study, a cohort of seventeen neurosurgical patients with left hemisphere brain tumours were recruited to investigate whether resection of certain white matter tracts affects the choice of hand selected for the execution of a goal-directed task (assembly of jigsaw puzzles). Patients performed the puzzles, but also tests for basic motor ability, selective attention and visuo-constructional ability, preoperatively and one month after surgery. Diffusion tractography of fronto-parietal tracts (the superior longitudinal fasciculus) and the corticospinal tract were performed, to evaluate whether resection of tracts was significantly associated with changes in hand selection. A complementary atlas-based disconnectome analysis was also conducted. Results showed a shift in hand selection despite the absence of any motor or cognitive deficits, which was significantly associated with patients with frontal and parietal resections, compared with those with resections in other lobes. In particular, this effect was significantly associated with the resection of dorsal fronto-parietal white matter connections, but not with the ventral fronto-parietal tract. Dorsal white matter pathways contribute bilaterally, with specific lateralised competencies, to control of goal-directed hand movements. We show that unilateral lesions, by unbalancing the cooperation of the two hemispheres, can alter the choice of hand selected to accomplish movements.

## Introduction

Handedness commonly refers to the tendency to use one hand over the other. Although the right and left hands of humans are nearly identical in their basic anatomy and motility, nearly 90% of the population show a strong preference for using the right hand to perform skilled movements (McManus, 2009; Corballis 2003). However, the subjective preference to select one hand to accomplish a specific task and the ability of this hand to do so are two related but not always corresponding dimensions of handedness (Bryden, Pryde & Roy, 2000; Herve et al. 2005; Angstmann et al. 2016). Significant scientific effort has been devoted so far to examining whether hand preference correlates with anatomical asymmetries (McManus et al. 2019), and how altering hand preference can affect neural structures (Marcori et al. 2019). Less attention has however been paid to evaluating whether hand preference can be altered as a consequence of changes in anatomical structure. This can be directly tested in the clinical setting by evaluating hand preference before and after neurosurgical interventions, which provides a unique opportunity to evaluate the neural basis of hand preference.

Manual dexterity primarily relies on the ability to perform independent finger movements, which requires monosynaptic corticospinal fibres from primary motor cortex to spinal motoneurons (Porter & Lemon, 1993). The corticospinal tract is broadly left-lateralised, with greater left to right decussation of the pyramids (Flechsig, 1876). Further, the left corticospinal tract has a more dorsal decussation at the midline in almost 90% of cases (Yakovlev & Rakic, 1966). While this was initially believed to be linked to right-handedness, both post-mortem and neuroimaging studies have demonstrated this to be unrelated to handedness (Lawrence & Kuypers, 1968; Kertesz & Geschwind, 1971; Westerhausen, 2007). A consistent finding has been that handedness is associated with morphology of the central sulcus, in proximity to the primary motor and somatosensory hand region (Amunts et al. 2000; Germann et al. 2019). However, this is not a rigid feature in that this region reshapes in corrected left-handers to follow a more ‘right-handed’ morphology, a consequence of an enforced shift in hand preference (Sun et al. 2012). It is well established that the precentral gyrus is highly plastic, thus handedness-related structural differences may reflect repeated lifelong use of one hand over the other (Steele & Zatorre, 2018; Simone et al. 2019). Given the lack of association between handedness and the asymmetry of cortical areas hosting corticospinal fibres for motor output, it is thus plausible that this difference may reflect structural asymmetry of pathways involved in earlier stages of action preparation.

Skilled manual action requires sensorimotor transformations to coordinate adequate muscle synergies to perform finger movements. Sensorimotor integration mainly requiring visual and somatic information, is mediated by a widespread fronto-parietal circuit (Turella & Lingnau, 2014), which has been well studied in macaques (Borra et al. 2017) but only partially in humans (Binkofski et al. 1999). Notably, neurons tuned to eye and hand movements in monkey fronto-parietal regions code primarily for the contralateral limb, but also for the ipsilateral limb (Cisek et al. 2003). This observation has also been demonstrated in humans, using transcortical magnetic stimulation (TMS) and functional magnetic resonance imaging (fMRI) (Schluter et al. 2001; Begliomini et al. 2008, Gallivan et al. 2013), indicating there is bilateral but left-lateralised specialisation for visuomotor control of movement that is handedness-independent (Sainburg et al. 2002; Begliomini et al. 2018). This is intriguing given the well-established right hemisphere dominance for visuospatial attention (Corbetta & Shulman, 2002). These results indicate that a bilateral frontal and parietal network mediates skilled manual actions, mainly involving the superior longitudinal fasciculus (SLF) running between the superior, middle and inferior frontal gyri and the superior and inferior parietal lobule (SLF I, II, III respectively) (Thiebaut de Schotten et al. 2011; Budisavljevic et al. 2017). In a previous study, we demonstrated that the structural asymmetry of these fronto-parietal tracts, rather than corticospinal asymmetry, differs between self-reported right- and left-handers, which is also linked with manual specialisation between hands on visuomotor tasks (Howells et al. 2018). Both groups had similar left fronto-parietal tract volume and performance with the right hand - the results were driven by differences in tract volume in the right hemisphere and left hand performance. This indicates that lateralised motor behaviour may not be the result of solely a more developed, efficient and thus dominant sensorimotor circuit in one hemisphere, but rather depends on the relationship between two homologous circuits in both hemispheres. Should this be the case, a lesion disrupting this cooperation may unbalance the system, resulting in an alteration of the motor behaviour of the hands. At present it is unknown whether motor behaviour is altered when this symmetry is disrupted by unilateral brain lesions, such as following neurosurgical procedures for the removal of a tumour.

Tractography is currently the only technique available for studying structural connections in the living human brain and is commonly used to evaluate the relationship between structural asymmetry and individual differences in behaviour (Catani et al. 2007; Forkel et al. 2014; Forkel & Catani 2018). In the clinical setting, tractography methods can estimate the extent of disconnection of specific white matter tracts by comparing the lesioned with the expected white matter anatomy known to be present in a healthy brain (Catani et al. 2012; Fox et al. 2018, Thiebaut de Schotten & Foulon, 2017). Surgical resection of tracts in one hemisphere alters their hemispheric asymmetry, thus changes in preferred hand use due to resection in specific regions may reveal neural structures that are relevant in mediating hand preference.

At present the most commonly used inventory scales to assess handedness lack the sensitivity to evaluate subtle changes in manual behaviour (Brown et al. 2006; Flindall & Gonzalez, 2018). These questionnaires measure the overall result of the hand selection process, but do not provide data to understand the underlying mechanisms themselves. Grasp-to-construct tasks are a useful means by which to evaluate lateralised motor behaviour in an ecological context, providing a quantitative measure of the interactions of each hand in both ipsilateral and contralateral space (Gonzalez et al. 2006, 2007). Putting together a jigsaw puzzle is therefore a useful way of testing hand selection to evaluate whether changes in lateralised manual behaviour following neurosurgical removal of brain tumours. The hand selected for the motor actions required during two phases of movement (reach-to-grasp and manipulation) during construction of puzzles was tested in seventeen patients in the preoperative phase and one month following the intervention. Patients performed basic motor tests to rule out any deficits in motor performance. As hand selection requires a significant cognitive load (Rosenbaum, 1980; Liang et al. 2018), we compared these results with performance changes on selective attention and visuoconstructional tasks to evaluate whether changes in hand selection were associated with deficits in these domains. Diffusion tractography of the main fronto-parietal tracts and the corticospinal tract were performed, to evaluate whether resection of specific tracts was associated with changes in hand selection after surgery. Based on the literature, we predicted that hand selection would be affected when resections occurred in white matter regions involving the branches of the fronto-parietal superior longitudinal fasciculus.

## Methods

### 2.1 Participants

Seventeen neuro-oncological patients who were candidates for awake surgery to remove a brain tumour were enrolled in this study. Patients were recruited using the following inclusion criteria: (i) a unilateral lesion in the left hemisphere, (ii) no previous surgery or radiotherapy (iii) no language or visual field deficits, (iv) no previous neurological or psychiatric conditions (v) no history of fractures involving the bones of the hand or fingers that might require restricted healing for longer than six months. All patients gave written informed consent to the surgical and direct electrical stimulation mapping procedure (IRB1299), and to the analysis of data for research purposes which followed the principles laid out in the Declaration of Helsinki. Patients were assessed for self-rated handedness using the Edinburgh Handedness Inventory (EHI, Oldfield, 1971). On this scale of hand preference patients could score between −100 and 100, where under −60 indicated completely left-handed, over 60 indicated completely right-handed and a score between −60 and 60 indicated mixed handedness.

**Table 1:**
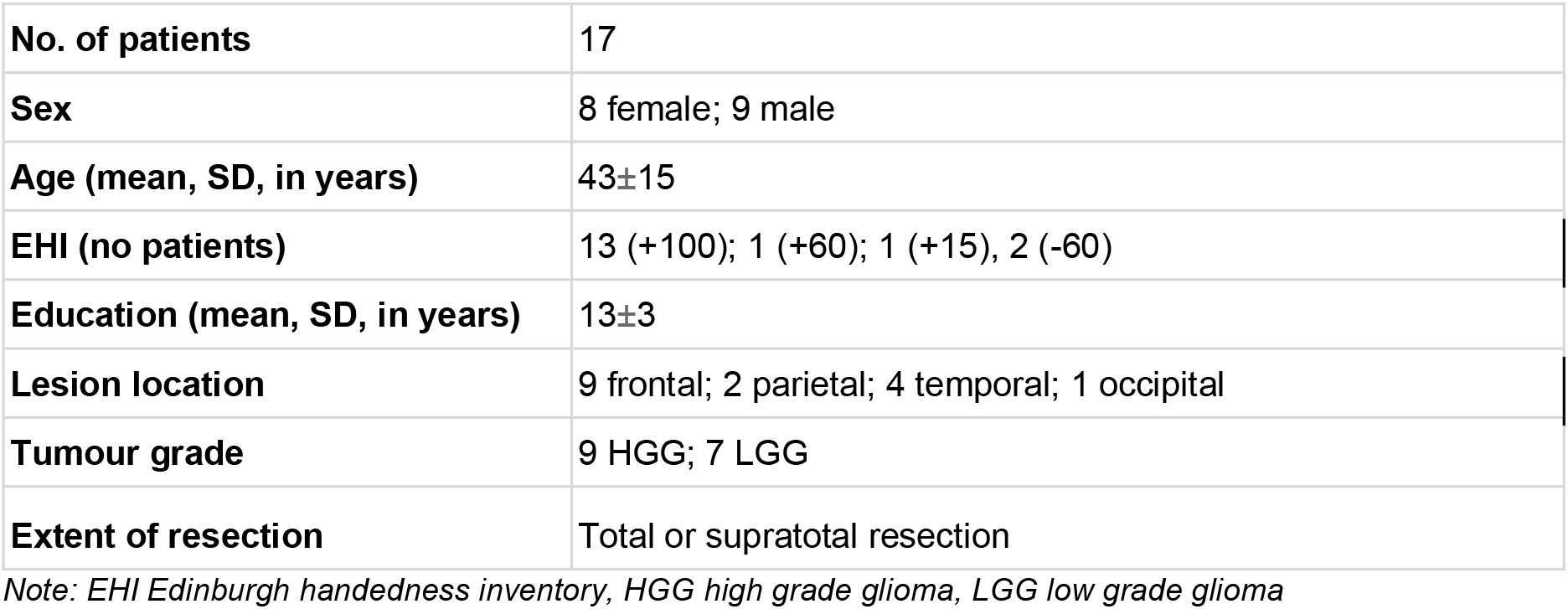
Demographic information

|  |  |
| --- | --- |
| **No. of patients** | 17 |
| **Sex** | 8 female; 9 male |
| **Age (mean, SD, in years)** | 43±15 |
| **EHI (no patients)** | 13 (+100); 1 (+60); 1 (+15), 2 (-60) |
| **Education (mean, SD, in years)** | 13±3 |
| **Lesion location** | 9 frontal; 2 parietal; 4 temporal; 1 occipital |
| **Tumour grade** | 9 HGG; 7 LGG |
| **Extent of resection** | Total or supratotal resection |

*Note: EHI Edinburgh handedness inventory, HGG high grade glioma, LGG low grade glioma*

### 2.2 Neuropsychological assessment

All patients underwent a comprehensive preoperative (1 week prior to surgery) and postoperative (1 month after surgery) neuropsychological assessment. This assessment included evaluation across cognitive domains including language, praxis, attention and executive function (for details see Puglisi et al. 2018). For the purpose of this study, and to exclude severe postoperative deficits that could affect the reliability of the postoperative assessment, we assessed changes between the pre- and postoperative timepoints for scores in visuospatial exploration (letter cancellation), visuoconstructional ability (Rey-Osterrieth Complex Figure), selective attention (Attentive Matrices) and auditory comprehension (Token Test).

#### 2.2.1 Assessment of hand performance

Hand performance was evaluated in two domains: arm-hand motor skills and praxis. The Action Research Arm Test (ARAT) is a simple test used to assess upper extremity movements with the dominant hand. It consists of 19 motor actions that are grouped into four subtests assessing four actions: grasp, grip, pinch, and gross movement. All items are rated from 0 (the movement is not possible) to 3 (normal performance of the task). The total score on the ARAT ranges from 0-57, with a higher score indicating better performance (Yozbatiran et al., 2008). We used 57 as the cut-off for this test. Patients without motor, sensory or visual deficits were assessed also for coordination and fine movement control using the Movement Imitation test for ideomotor apraxia (De Renzi, 1980). It consists of twenty-four gestures of different complexity that are imitated by the patient, requiring independent movement of the hands. Each imitation trial is rated from 0 (impossible to replicate the movement) to 3 (correct imitation at first presentation). The total score ranges from 0-72, where a score of 52 is the cut-off for normal performance.

#### 2.2.2 Assessment of hand preference: jigsaw puzzle task

During the neuropsychological assessment and while comfortably seated in front of a table, patients were asked to assemble two different jigsaw puzzles to evaluate spontaneous hand preference in a ‘naturalised’ setting (Gonzales et al. 2006). Each puzzle was of a standard size (17cm x 17cm) and made up of 25 equally sized pieces (Supplementary Video). The underside of the pieces were labelled with ‘1’ or ‘2’ to indicate the hemispace in which they were to be presented. The pieces were distributed across each side of the tabletop with the same number of pieces on each side. The patient was seated exactly facing the middle of this distribution and provided with a central puzzle piece directly in front of them, for orientation. An image of the completed puzzle image was displayed opposite the patient for reference. The patients were asked to place each hand on the table face down and then to reproduce the puzzle as fast and as accurately as possible and were blinded to the purpose of the study (no instruction was given as to which hand to use). Patients were asked to take one piece at a time and replace it if they could not fit it into the puzzle. The patients’ hands were video recorded by a camera position directly in front of the patient, tilted downward to provide a view of the action of both hands. Patients were given three minutes to complete each puzzle and then asked to stop, even if the puzzle was not completed. The order of presentation of each puzzle was counterbalanced between patients.

Performance on the two puzzles were scored offline using the video-recordings, by two neuropsychologists blind to whether performed pre- or post-operatively (see Supplementary Video). The performance was evaluated in two action phases: reach-to-grasp and manipulation. First, each video was analysed to record the hand used every time a piece of the puzzle was reached for and grasped (e.g.; Figure 1a). It was also recorded whether the hand used was reaching to grasp a piece within its hemispace (e.g. right hand within right hemispace, R, left hand within left hemispace, L) or whether it reached to grasp within the opposite hemispace (e.g. right hand into left hemispace, Rx; left hand within right hemispace, Lx). As the effort required to reach across hemispace was higher, the last condition was given a higher weight (Elliott et al. 1993; Liang et al. 2018). We used the average across the two trials to create a final score of lateralised hand selection, calculating a lateralisation index calculated as (R + (1.5 x Rx)) or (L + (1.5 x Lx)). A score of −1 reflects selection solely of the left hand, a score of +1 reflects selection solely of the right, while a score of 0 reflects selection of both hands equally. When one hand would reach and grasp a puzzle piece, this was sometimes passed to the other hand for positioning. Thus, each video was also scored for the hand that rotated the puzzle piece into the appropriate configuration and then fit it into position (Figure 1b). This was a cooperative movement as the other hand generally played as a supportive role, by holding the puzzle in place. The final score for each hand was calculated based on the total number of manipulations performed by each hand and a similar lateralisation index of hand selection was created (average across the two trials). For each of the two scores, the proportion of right hand use out of the total grasps or manipulations was also calculated (R/(R+L)).

**Figure 1.**
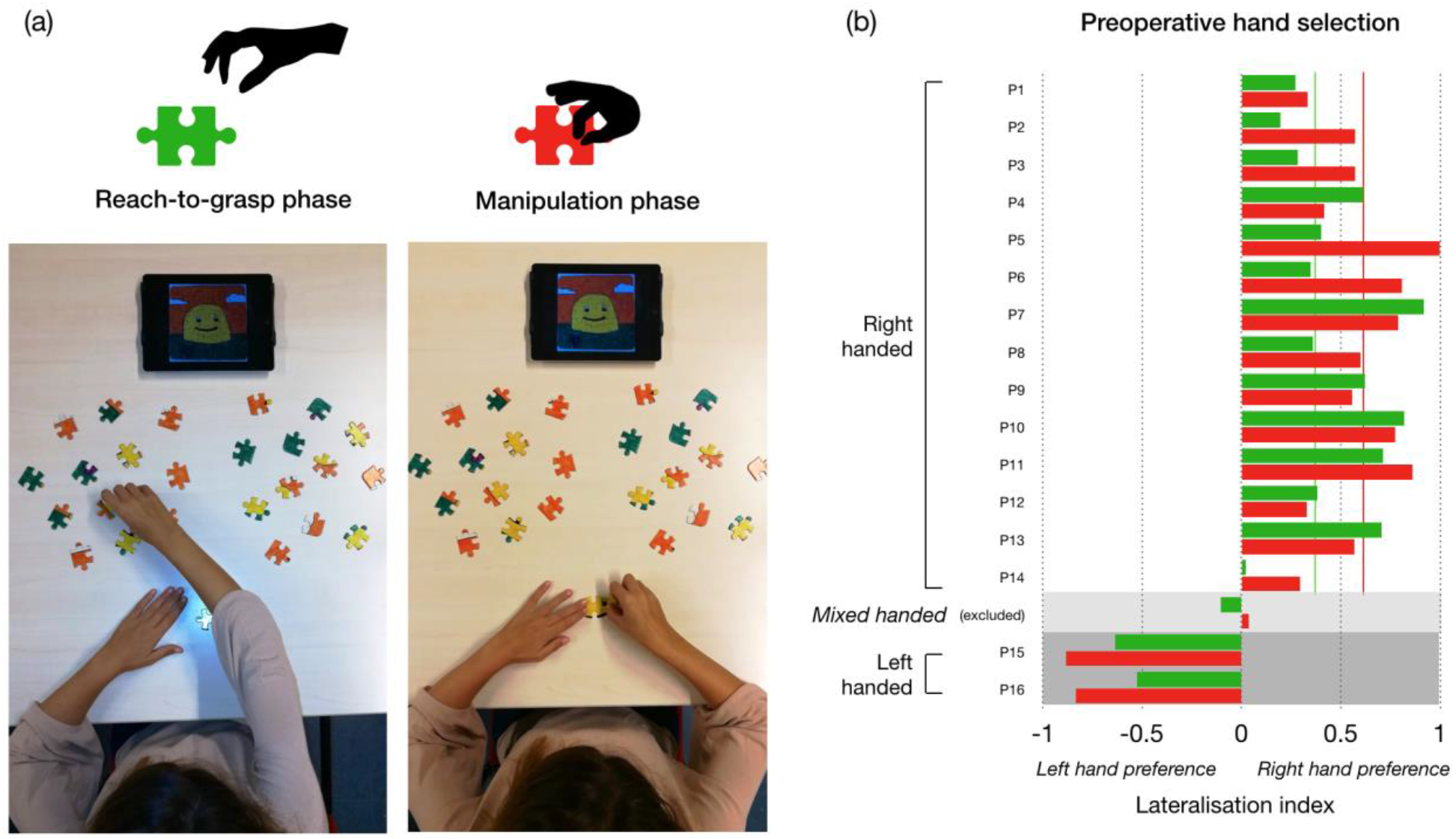
Photographs showing layout of puzzle task and hand movement during the (a) reach-to-grasp (green) and (b) manipulation (red) phases (see also Supplementary Video). (c) A bar graph shows hand selection in the preoperative time point for both phases of the puzzle task (lines in red and green reflect median score for right-handed population). The direction of hand use is consistent with hand preference reported on the Edinburgh Handedness Inventory.

#### 2.2.3 Statistical analysis of neuropsychology data

Multiple paired t-tests were used to assess differences between pre- and post-operative scores related to: a) hand motor function, b) hand selection for the puzzle task and c) cognitive status. Multiple repeated measures mixed ANOVAs were employed to assess the interaction between clinical or demographic variables and hand selection in the pre- and post-operative timepoints. We used timepoint (pre vs post) as the within-subjects factor, resected lobe, education and sex as between-subjects factors, and resection volume and age as covariates. As we hypothesised that fronto-parietal resections would have a significant impact on hand selection, for the resected lobe factor we categorised patients into two groups: those with resections predominantly in the frontal or parietal lobe, and those with resections in the temporal or occipital lobe.

### 2.3 Neuroimaging acquisition

All patients underwent a clinical MR imaging sequence one day before surgery, and at the one-month follow-up. Preoperative MRI imaging was performed on a Philips Intera 3T scanner (Koninklijke Philips N.V. Amsterdam, Netherlands), and acquired for characterisation of lesion morphology and volume. A post-contrast gadolinium T1-MPRAGE sequence was performed using the following parameters TE:2.7ms, TR:95.4s, FOV: 176 slices, slice thickness: 1mm and a T2-FLAIR, as part of the clinical routine.

Nine patients also underwent a High Angular Resolution Diffusion Imaging (HARDI) sequence for clinical purposes using an 8-channel head coil. A spin echo, single shot EPI sequence was performed with 73 directions collected using a b-value of 2000s/mm3, and seven interleaved non-diffusion weighted (b0) volumes (TE:96ms, TR 10.4ms). The acquisition had a matrix size of 128×128 with an isotropic voxel size of 2mm3.

#### 2.3.1 Neuroimaging preprocessing and analysis

Volumetric analysis was used to define tumour volume using BrainLab software (Smartbrush). Resection cavities were delineated on the postoperative T1 and registered to a preoperative diffusion-weighted imaging map (Anisotropic Power, Dell’Acqua & Tournier, 2018) using the Clinical Toolbox in SPM (Rorden et al., 2012).

Diffusion imaging data was visually inspected for outliers, corrected for signal drift, reordered and corrected for head motion and eddy current distortions using ExploreDTI (www.exploredti.com, Leemans et al. 2009). Standard diffusion tensor models cannot show multiple fibre orientations within a voxel therefore are not suitable for evaluating fronto-parietal tracts (Thiebaut de Schotten et al. 2011). We used an advanced algorithm based on spherical deconvolution to model the orientation distribution function, using a damped Richardson-Lucy algorithm (Dell’Acqua et al. 2010). The following settings were used: ALFA = 1.7, 300 iterations, n= 0.001, v = 8, and an absolute threshold of 0.001. Whole brain tractography was calculated using a step size of 1mm, with a constraint to display streamlines between 15 and 200mm in length. Euler interpolation was used to track streamlines using an angle threshold of 45 degrees. All spherical deconvolution modelling and whole brain deterministic tractography was performed using StarTrack software (Dell’Acqua et al. 2012; www.mr-startrack.com).

#### 2.3.2 Tractography dissections

Virtual dissections of the three branches of the superior longitudinal fasciculus (SLFI-III) and the precentral component of the corticospinal tract were performed using a ROI-based approach. The regions of interest used as segment the SLF I-III are described in detail in Thiebaut de Schotten et al. 2011 and Howells et al. 2018. The dorsal branch of the SLF (SLF I) connects the superior parietal lobule with the superior frontal gyrus, running anterior and parallel to the cingulum but distinct, separated by the cingulate sulcus (Thiebaut de Schotten et al. 2012). The middle branch (SLF II) connects the posterior inferior parietal lobule (angular gyrus) with the middle frontal gyrus including the frontal eye fields. The ventral branch (SLF III) connects the inferior frontal gyrus and ventral precentral gyrus with the anterior inferior parietal lobule (supramarginal gyrus) and intraparietal sulcus. For the purpose of this study, the corticospinal tract was defined as the streamlines extending from the precentral gyrus to the brainstem (Catani & Thiebaut de Schotten 2016). The postoperative MR with the delineated resection cavity was registered and overlaid on the preoperative diffusion tractography using a similar approach to that described in Puglisi et al. (2019). By using the resection cavity as an inclusion ROI, we could estimate the percentage of streamlines that were disconnected by the resected region.

#### 2.3.3 Estimation of tract resection

We used a supplementary approach to estimate the disconnection of tracts in the entire cohort. While use of atlas-based tract estimation tools is challenging in patients with tumours that may displace or disconnect tracts, it is a useful adjunctive tool to estimate tract-based lesion-symptom associations, to complement findings identified with tractography. We used the online platform “Megatrack”, a HARDI-based tractography atlas and lesion tool (https://megatrackatlas.org, Stones et al. OHBM 2019), to estimate the extent of disconnection of white matter tracts, based on the percentage of disconnected streamlines as a proportion of the total making up the fibre bundle. Although a number of tract atlases are available (Rojkova et al. 2016), this approach is highly relevant in this case, as one can take specific demographic factors can be taken into account when predicting likely tract volume such as handedness (Howells et al. 2018).

A mixed repeated measures ANOVA was used to evaluate whether there was an interaction between resection of a tract and a change in neuropsychological performance across tests (puzzle - reach-to-grasp or manipulation phase, Rey figure, Attentive Matrices). We used timepoint (pre vs postsurgery) as a within-subjects factor and the status of each of the three branches of the superior longitudinal fasciculus (resected/preserved) as a between-subjects factor.

## Results

### 3.1 Assessment of motor and cognitive abilities

#### Upper limb motor skills

Motor assessment was performed to exclude alteration of motor ability of the dominant hand before and after surgery. The ARAT was used to test the ability of the dominant hand in performing four basic motor actions (i.e. grasp, grip, pinch, and gross movement). The task was fully accomplished (i.e. all actions were performed with full scores) by all patients at both timepoints (Figure 2).

**Figure 2.**
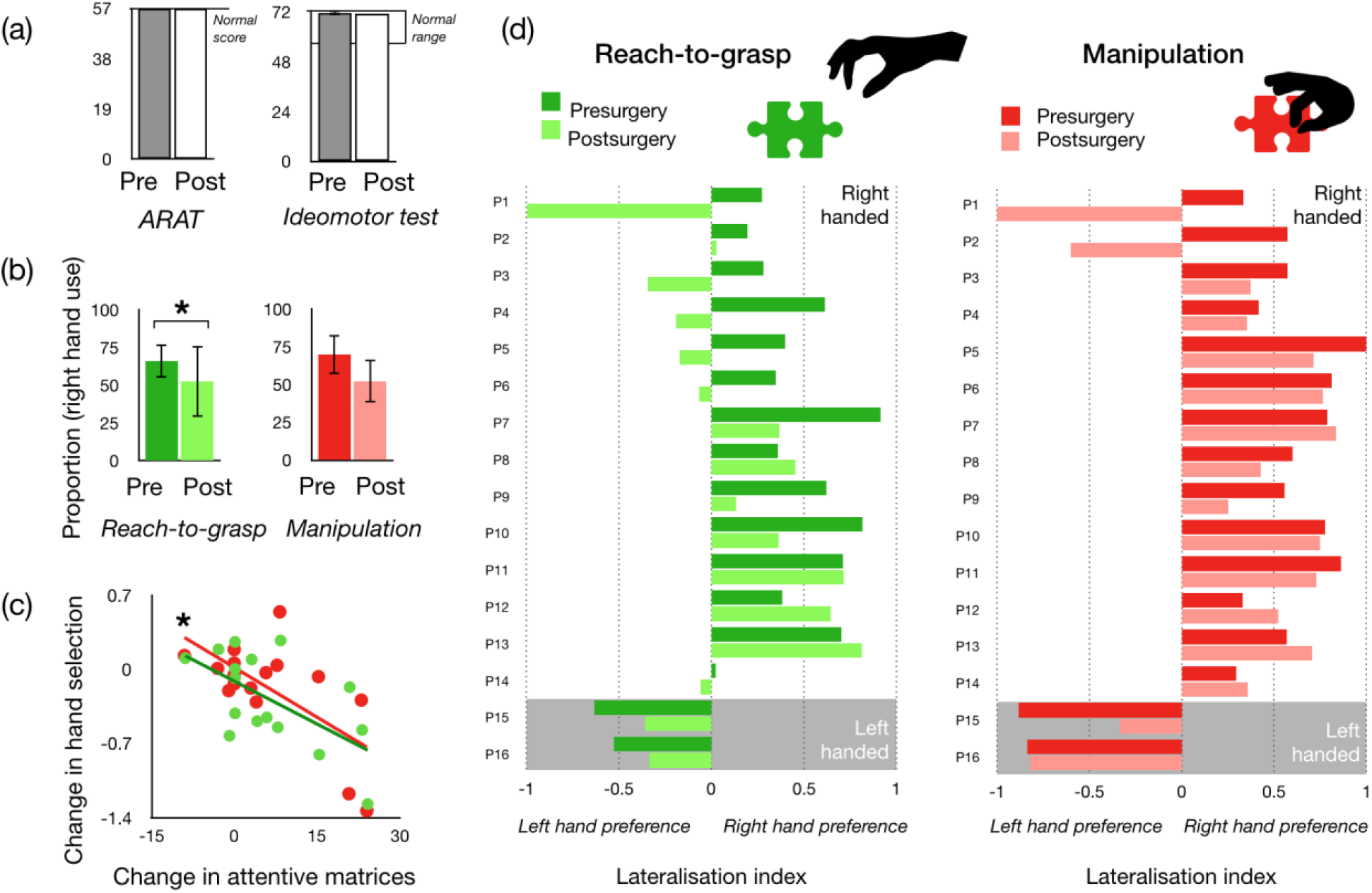
Changes in neuropsychological scores before and after surgery. (a) Motor ability before and after surgery. (b) The change in hand selection in the two phases of the puzzle. (c) A scatter graph showing the significant association between change in hand selection for each phase and change in score on the attentive matrices. A negative score indicates a shift to non-dominant hand use, or an improvement in the selective attention score. (d) the individual scores for each patient are shown in the pre- and post-operative phases. Note: * reflects significance level of p<0.05.

#### Praxis ability

There was no significant change in score on the ideomotor test (t(16)=126, p=0.126), indicating no praxis deficits were evident before or after surgery.

#### Language comprehension

No patients experienced persistent postoperative aphasia, and their performance on the Token test for auditory comprehension, despite a decrease (t(15)=2.7, p=0.014), was within the range of normality in the postoperative phase (cut-off 22.5). All patients were therefore able to understand the instructions given for the task.

#### Attentional processing

In line with the postoperative clinical course, a slight reduction in cognitive performance was observed in selective attention (t(15)=2.5, p=0.023). The difference in omitted letters between right and left hemifields in the cancellation test was assessed in the pre- and postoperative phases. There was no significant change in visual field exploration between the two timepoints (t(15)=−1.0, p=0.3). None of the patients showed hemispatial neglect.

No patients experienced any postoperative sensory deficits. One patient presented with hemianopia in the first month follow-up, which fully recovered subsequently (Patient 1). In order to assess any potential postoperative difficulties in task completion we compared the number of correct pieces placed at the end of the puzzle task between the two timepoints. The mean number of pieces correctly placed was 14.9 out of 25 (s.d. 7.4) in the preoperative time point and 14.7 out of 25 (s.d. 7.3) in the postoperative time point. A paired samples t-test showed no significant difference between time points (t(15)=−0.249, p=0.8).

### 3.2 Assessment of hand selection

Patients were asked to complete a self-rated handedness inventory (Edinburgh Handedness Inventory, EHI) before and one month after surgery. Fourteen patients were right-handed (+60 on EHI), two patients were left-handed (−60 in EHI) and one patient was mixed handed (+ 37.5 on EHI). No patients reported any change in the EHI score in the one month follow up.

#### Assessment of task consistency before and after surgery

A comparison of hand selection, as measured by the lateralisation index, was conducted between trials (first vs second puzzle). Lateralised hand selection was highly correlated between the two trials in the preoperative (r=0.8, p<0.001) and postoperative phase (r=0.6, p<0.003), indicating the test had good intraindividual consistency for assessing hand selection in both reach-to-grasp and manipulation phases.

#### Assessment of consistency of hand selection in the two phases of the puzzle task

Hand selection on the puzzle was compared with the patient’s self-reported hand preference in the preoperative time point. All patients used their dominant hand more than the non-dominant hand for both phases of the puzzle (reach-to-grasp and manipulation; Figure 1c), with the exception of the mixed handed patient who showed an inconsistent hand preference. This patient was excluded from subsequent neuropsychological analysis. We evaluated the consistency in the hand selected for both reach-to-grasp and manipulation phases. A bivariate analysis showed a strong correlation between the two phases of the puzzle task in the preoperative phases (r2=0.8, p<0.001). An ANOVA showed no effect of sex on lateralised preoperative hand selection in either phase (reach-to-grasp: F(1,14)=0.227, p=0.6; manipulation: F(1,14)=0.37, p=0.6).

**Table 2:**
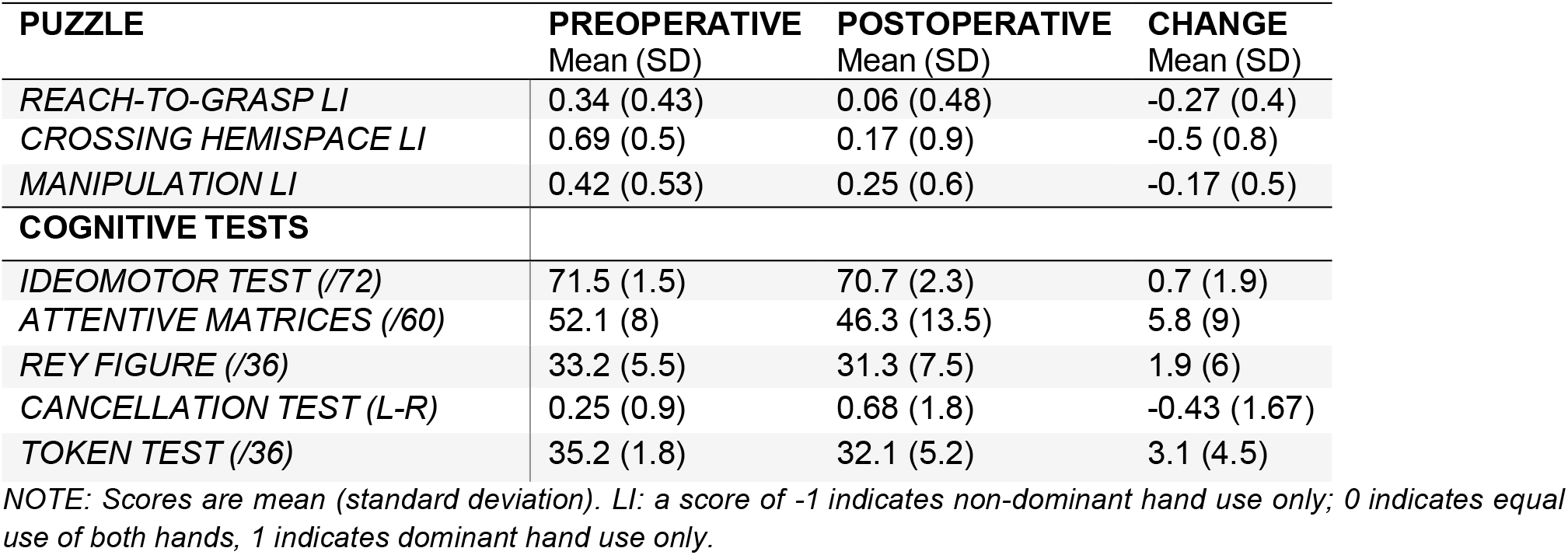
Neuropsychological scores before and after surgery

#### Assessment of hand selection before and after surgery

Multiple repeated measures ANOVAs were used to assess the interaction between clinical or demographic variables and the change in hand selection before and after surgery. The ANOVAs revealed no significant interaction between change in hand selection and education, sex, resection volume or age. The only significant interaction was between resected lobe and hand selection for both reach-to-grasp (F(1,14)=6.87, p=0.02) and manipulation phases (F(1,14)=5.06, p=0.04). A significant difference in hand selection before and after surgery emerged in patients with frontal or parietal resections, but not when resection affected the temporal or occipital lobes (Figure 3a).

**Figure 3.**
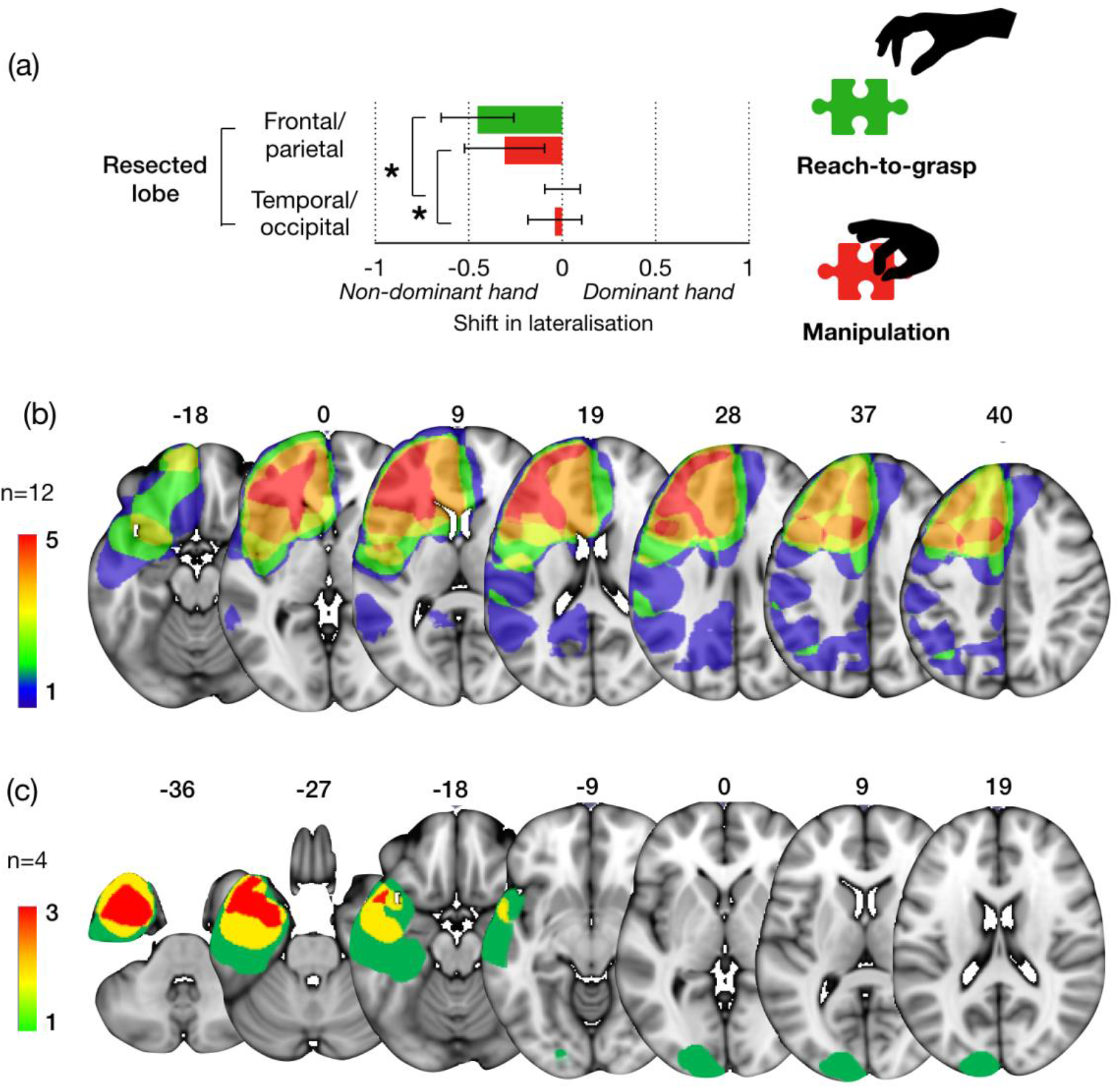
(a) Bar graph showing the shift in hand selection by resection group (b) Anatomical distribution of resections within the frontal and parietal lobe. The lesion location of the mixed handed patient is not included here (c) Anatomical distribution of resections within the temporal and occipital lobe. *p<0.05.

We finally compared cognitive scores with hand selection on the puzzle task. Bivariate analysis showed a significant association between change in selective attention performance and hand selection for reach-to-grasp (r2=0.605, p=0.01) and manipulation (r2= 6.01; p=0.014; Figure 2). The greater shifts toward non-dominant hand use were correlated with lower scores on the selective attention test. No significant correlations between change in visuoconstructional ability or auditory comprehension, and change in hand selection for reach-to-grasp or manipulation were observed.

### 3.3 Effect of resected region on hand selection

Awake neurosurgery was performed in all patients, with the aid of the brain mapping technique, using functional borders to achieve total or supratotal resection for tumours distributed across the left hemisphere. All regions of the precentral gyrus for which motor evoked potentials of the hand could be evoked by direct electrical stimulation were preserved in all cases (Bello et al. 2014; Figure 3). Further, a new tool designed to assess and preserve eloquent regions controlling complex non-visually guided hand actions was used during awake brain mapping in these patients (see previous studies – Fornia et al. 2019). Mean resection volume was 76.7ml (s.d. 71.3).

**Table 3.** Tractography measurements in right and left hemisphere

|  | TRACT MEASUREMENTS (L) | TRACT MEASUREMENTS (R) | TRACTOGRAPHY DISCONNECTION IN LEFT HEMISPHERE |
| --- | --- | --- | --- |
| **TRACTS** | Mean Volume (SD) | Mean Volume (SD) |  |
| SLF I | 11473.5 (3439) | 14444.25 (2532) | 5/9 cases |
| SLF II | 13373.25 (8247) | 13166.875 (5873) | 4/9 cases |
| SLF III | 7722.4 (3144) | 12033.5 (3349) | 4/9 cases |
| CST | 11455.4 (2431) | 9528.6 (1266) | 0/9 cases |

NOTE: SLF: superior longitudinal fasciculus, CST corticospinal tract

The tractography results showed that, following surgery, the dorsal branch of the superior longitudinal fasciculus (SLF I) was disconnected in 5/9 cases, the middle branch (SLF II) in 4/9 cases and the ventral branch (SLF III) in 4/9 cases. The corticospinal tract was intact in all patients. We examined changes in hand selection in the two phases of the puzzle task (reach-to-grasp and manipulation) between patients with specific branches of the superior longitudinal fasciculus resected or preserved. We observed a trend to show greater shift in hand selection toward non-dominant hand use following resection of the SLF I or SLF II (Figure 4b). No consistent result was associated with resection of the SLF III.

**Figure 4.**
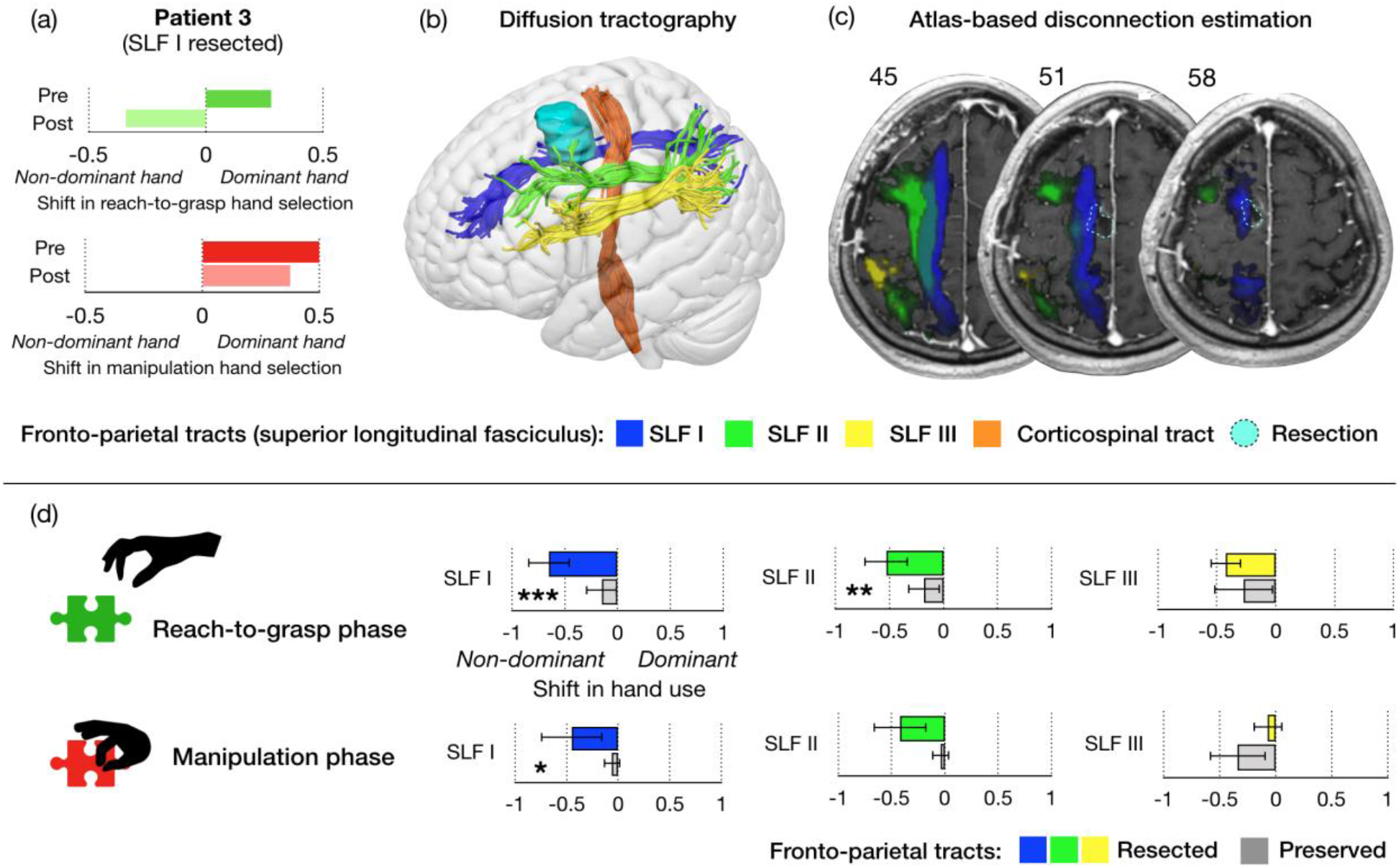
Example of one patient included in the study (patient 3), (a) demonstrating change in hand selection on different phases of puzzle. (b) Preoperative diffusion tractography dissections of the four tracts in this patient are shown with an overlay of the resection cavity (cyan), showing the SLF I was resected (c) Megatrack atlas-based estimation of white matter disconnection shown on the postoperative T1 also indicated complete resection (d) Boxplots showing group differences in change in hand selection related to resection of specific tracts in the different phases of puzzle performance. ***p<0.001, **p<0.01, *p<0.05

As under half of patients underwent diffusion tractography (9/16) patients, we confirmed our results using an atlas-based approach. Again, this approach confirmed the preservation of all precentral projections of the corticospinal tract. The dorsal fronto-parietal tract (SLF I) was resected in 6/16 patients, the middle branch (SLF II) in 8/16 patients and the ventral branch (SLF III) in 7/16 patients. There was good correspondence between the atlas-based and tractography-based approach.

Multiple repeated measures ANOVA showed an interaction between (atlas-based) tract resection and shift in hand selection on both reaching and manipulation phases (Figure 4b). Patients with the left dorsal fronto-parietal branch (SLF I) resected showed a significantly greater shift toward non-dominant hand use in the reach-to-grasp phase compared to patients submitted to a resection preserving the same tract (F(1,14)=16.45, p=0.001). This same results were observed for the manipulation phase (F(1,14)=4.4, p=0.05). Resection of the SLF II resulted in a shift in hand selection in the reach-to-grasp (F(1,14)=9.9, p=0.007) but not the manipulation phase. Resection or preservation of the SLF III did not affect the hand selection in the task. Overall the analysis point to the left and middle fronto-parietal branches as significant tracts involved in hand selection.

Additionally, we evaluated whether resection of these tracts was related to a change in cognitive performance on the selective attention and visuoconstructional tasks. Resection of the SLF I and SLF II result in a significant shift in hand selection during the task, and patients with resection of the SLF I also showed a trend toward higher incidence of deficits on the selective attention task at 1 month following surgery (F(1,14)=4.3, p=0.058), as did those with resection of the SLF II (F(1,14)=4.3, p=0.056). No associations between resection of these tracts and visuoconstructional task performance before and after surgery was observed.

## Discussion

In neurosurgical patients with left hemisphere brain tumours, we investigated whether resection of fronto-parietal white matter pathways was associated with a shift in hand selection, assessed using a puzzle assembly task. Our results show that subtle changes in hand selection occurred following frontal and parietal resections, despite no primary deficits in motor ability. Patients primarily selected the self-reported dominant hand (based on the EHI) for both reach-to-grasp and manipulation phases of puzzle assembly, however there was a shift toward the increased use of their non-dominant hand for this task in the postoperative phase. This hand selection shift was significantly correlated with the surgical resection of superior and middle fronto-parietal white matter connections (i.e. SLF I and II), but not the inferior fronto-parietal branch (SLF III). Our results suggest that the relationship between brain structure and lateralised hand motor behaviour is reciprocal: forced alteration of spontaneous manual preference can affect structural hemispheric asymmetries (Sun et al. 2012), but also lesions altering brain structure can produce subtle shifts in lateralised motor behaviour.

Everyday interactions require complex highly skilled hand movements which are either performed unimanually, or more commonly, requiring bimanual cooperation. The preference to use one hand over the other to perform complex motor tasks is a distinct feature of our species, a lateralised behaviour referred to as handedness. Hand-object interaction requires independent finger movements to be orchestrated based on the properties of the object and the goal of the action. Thus, lateralised hand use is unlikely to depend solely on asymmetry of neural structures in change of final motor output, such as the dimensions of the precentral gyrus or corticospinal tract. A considerable body of work has indicated that cooperative interplay of both hemispheres is required for movement, and further that each hemisphere is responsible for different aspects of motor programming for complex actions (Sainburg et al. 2002). When considering grasping and hand-object manipulation, fronto-parietal connections are essential in providing the motor program with visual and somatosensory information required to achieve adequate hand shaping and control in both monkeys and humans (Borra et al. 2017; Turella & Lingnau 2014). Three parallel branches of the superior longitudinal fasciculus convey sensorimotor transformations between frontal and parietal regions, each of which has different patterns of structural asymmetry (Thiebaut de Schotten et al. 2011). This interhemispheric asymmetry has been associated with specific aspects of upper limb kinematics in healthy adults, precisely for different phases of visuomotor processing needed for reach-to-grasp movements (Budisavljevic et al. 2016) and may have a genetic basis (Wiberg et al. 2019). We recently demonstrated further that hemispheric asymmetry of the dorsal fronto-parietal tract differs between self-reported right- and left-handers, with a greater left-lateralisation in right-handers, and right-lateralisation in left-handers (Howells et al. 2018). Asymmetry of these fibres was also associated with manual specialisation between the hands, measured using relative unimanual performance between hands on a pegboard task. This indicates that relative contributions from both hemispheres are involved in facilitating task performance with each hand.

Taken together, recent evidence indicates that lateralised motor behaviour, whether relating to hand selection or manual ability, is linked to the interplay of both hemispheres, each in charge of specific aspects of motor programming. In line with this hypothesis, unilateral lesions should result in alteration of lateralised motor behaviour related to a specific feature of motor programming by unbalancing interhemispheric interplay dependent on certain structural asymmetries. The neurosurgical setting thus provides an opportunity to observe the consequence of selective lesions. Our results indicate that neurosurgical resection of both frontal and parietal left hemisphere regions alters motor behaviour, shifting hand selection toward increased non-dominant hand use one month after the procedure. In particular, this was related to resection of those regions connected by the dorsal and middle branch of the superior longitudinal fasciculus (SLF I and II).

### 4.1 Fronto-parietal resection affects hand selection but not motor ability

A key result that emerged from our study is that the neurosurgical resections performed in premotor and parietal regions did not impair gross motor skills of the dominant hand, as all patients performed within the normal range on both basic motor and ideomotor apraxia tests (Figure 2). Preservation of these functions was due to the intraoperative cortical and subcortical electrical stimulation awake mapping procedure, used to identify eloquent structures during resection and hence producing functional borders to resection (Bello et al. 2014). In this case, patients use a dedicated object manipulation tool demonstrated to preserve praxis function (Rossi et al. 2018; Vigano et al. 2019; Fornia et al. 2019). Resection of fronto-parietal tracts in the left hemisphere did not impair motor ability itself, but rather caused a shift (and rarely a flip) in hand selection for reach-to-grasp movements: the dominant hand was still used primarily over the non-dominant hand in the postoperative timepoint, although to a lesser extent. This indicates that presurgical hand preference was still preserved, however the strength of its dominance over the other hand decreased. A prominent model of bilateral hemispheric interplay in control of hand movement indicates that the left hemisphere (in right-handers) is specialised for predictive control of limb dynamics, whereas the right hemisphere is specialised for impedance control and positional stability in unanticipated perturbations (Sainburg, 2002). Both hemispheres contribute to the motor program with different competencies. Damage to the left hemisphere may thus interrupt the ballistic component or timing of movements which may affect the trajectory of the dominant hand. The hand selected for the task may therefore change to compensate and to ensure the goal of the task is still achieved.

Notably, our results also showed that there was a similar shift toward right-hand use in the left-handers tested, following the left hemisphere resections. This may provide preliminary evidence to support the hypothesis that the left hemisphere is specialised for visually guided dominant hand grasping, in both left and right-handers (Begliomini et al. 2018). Altogether this data supports the hypothesis that bilateral fronto-parietal tracts support complex hand movements, for which the balance of communication between hemispheres may support hand selection for goal directed actions (Budisavljevic et al. 2016; Howells et al. 2018). However, a second point arises from these results: as goal-directed movements could still be performed, the inclination to use the non-dominant, ipsilesional hand more (or dominant hand, less) may reasonably be related to computations reflecting the influence of a higher cognitive mechanism such as movement intentionality or executive function.

### 4.2 Hand preference and online control of movement

The dorsal fronto-parietal branch (SLF I) extends between superior frontal and anterior cingulate cortices, and the precuneus and superior parietal lobule and has been traced in both monkeys and humans (Petrides & Pandya, 1984; Thiebaut de Schotten et al. 2012). Despite running parallel to the cingulum, post-mortem studies have demonstrated that it is a distinct tract, separated by the cingulate sulcus (Yagmurlu et al. 2016; Komaitis et al. 2019). The frontal terminations of the SLF I include the preSMA, which codes for both contralateral and ipsilateral limb movements (Gallivan et al. 2013) and plays a critical role in translating higher level goals to action (Wang et al. 2019). Other cortical regions in the superior frontal gyrus including the frontal eye fields play an important role in attention and working memory (Boisgueheneuc et al. 2006). The SLF I connects all of these regions with the superior parietal lobule, crucial for orienting actions within space, using visual information to code target location and movement direction, transforming spatial targets into movement vectors (Goodale & Milner, 2018; Barany et al. 2014; Gallivan & Culham, 2015). The superior parietal lobule can directly influence motor output through M1, but also is connected with premotor cortex to form major relays for coordinating reach-related grasping movements (Monaco et al. 2011; Cattaneo et al. 2019). Notably, the function of the superior parietal lobule relates to online monitoring of one’s own body - lesions in this region can cause disorders of self-awareness such as fading limb, alien hand or autotopagnosia (Wolpert et al. 1998; Herbet et al. 2019). The bilateral SLF I likely conveys neural impulses for online control of movement of both hands, and our results show that disconnecting this tract in the left hemisphere causes a shift toward non-dominant hand use when exploring peripersonal space. In a previous study, we reported handedness-related differences in hemispheric asymmetry of SLF I volume in healthy adults, a measurement likely reflecting enhanced speed of conduction (Howells et al. 2018; Drobyshevsky et al. 2005). Further, damage in the right hemisphere causes hyperexcitability of parieto-motor connections in the left fronto-parietal network (Koch et al, 2008). Considering this evidence, one hypothesis may therefore be that hand selection, as measured by our test, is a reflection of top-down online monitoring of one hand, due to faster and more efficient movement intentionality. Thus, a shift in hand selection may reflect lower ‘power’ of the dominant hand in this regard, or conversely, an upregulation of movement intention in the non-dominant hand that disturbs the other. Further investigation is however required to test this theory.

### 4.3 Hand preference and attentional processing

A recent combined magnetoencephalography-tractography study has also linked differences in structural asymmetry of the SLF I to selective attentional processes, measured in synchronisation of alpha and gamma band oscillations (Rhys Marshall et al. 2015). While the role of selective attention in action selection has been well described (Castiello, 1999), our results further show an association between selective attention and dominant or non-dominant hand selection. Patients with greater shift toward non-dominant hand use following surgery also had reduced selective attention ability, despite no impairment in visual search strategies in either hemispace. Further, our results also show that resection of the second branch of the SLF (SLF II) connecting the middle frontal gyrus with posterior inferior parietal lobule (the angular gyrus) was associated with changes in hand selection, with a similar trend for selective attention. Importantly, this tract connects neural regions within two important attention networks: the dorsal attention network (DAN; SLF I) and the ventral attention network (VAN; SLF III) (Corbetta & Shulman, 2002). Individual differences in structural asymmetry of the SLF II are associated with attentional biases in healthy adults, detected using behavioural tasks such as the line bisection task (Thiebaut de Schotten et al. 2011). Recent TMS-tractography combined studies have also demonstrated this tract also plays a key role in online monitoring and movement correction of actions (Koch et al. 2010; Rodrigues-Herreros et al. 2015). Thus, the SLF I and SLF II are likely to both be involved in top-down attentional processing as well as mediating online control of movement. Supporting this, there was a trend toward patients with resection of the SLF I and/or SLF II having greater declines in performance on the selective attention task in the postoperative phase. This suggests there may be a link between attentional processing and lateralised hand selection. Focusing on goal-relevant stimuli while ignoring distractors requires executive control to efficiently allocate attentional resources, which is theorised to be supramodal (Lavie et al. 2005; Spagna et al. 2015; Ptak et al. 2017). In 1980, Rosenbaum investigated reaction time for reaching, altering the pre-cues such as direction, distance and the hand to be used for the movement. He demonstrated that reaction time was reduced most substantially when hand selection was cued, indicating this decision-making process has a considerable cognitive load. Executive control of attention therefore may extend also to allocating motor attention toward selection of one hand over another (Rushworth et al. 2003).

### 4.4 Limitations

Resection of the ventral fronto-parietal branch (SLF III) connecting the inferior frontal gyrus and ventral precentral gyrus with the anterior inferior parietal lobule did not seem to affect hand selection in our patient cohort. Given that structural asymmetry of this tract has been associated with both kinematics of reach-to-grasp movements and handedness, this result was unexpected (Budisavljevic et al. 2016; Howells et al. 2018; Wiberg et al. 2019). A possible explanation might be that the paradigm used to construct a puzzle may be more adequate to test online control of movement in peripersonal space, rather than specific hand shaping for grasping, therefore it may not have been sensitive enough to detect subtle changes in skilled motor actions.

Studying the relationship between clinical manifestations and lesions in patients with brain tumours is of great aid in that, unlike in situations of vascular insult, lesions are constrained and more focal, and it is possible to assess neuropsychological performance before as well as after a neurosurgical intervention. However, there are a number of limitations that deserve discussion. First, it is challenging to assess whether or to what extent brain function is impaired in areas of diffuse tumour infiltration. In this study, the growth of a tumour may already have affected hand preference, which may explain why right-hand preference was not as strong as expected based on previous studies (90% right-hand grasps in right-handers e.g. in Gonzalez et al. 2006). Further, brain tumours are a rare disease, thus the patient cohort tested was relatively small. With a larger patient cohort, we would have been able to conduct voxel-lesion symptom mapping and more sophisticated statistical analyses that would be better able to confirm our preliminary results (e.g. Foulon et al. 2018). Moreover, our evaluation of motor ability was relatively crude, and kinematic analysis would better be able to rule out the impact of subtle motor impairments and their effect on hand selection.

### 4.5 Conclusions

Handedness likely consists of a number of dimensions, each of which underlie lateralised motor behaviour for a circumscribed set of tasks. Given that handedness does not have a one-to-one relationship with manual specialisation, the different items and skills required for different tasks designed to investigate this topic may yield different insights into preferred use of one hand for interaction with the immediate environment (Todor & Doane, 1977). We here confined the investigation of hand preference to a task involving completion of a jigsaw puzzle - requiring reaching to grasp pieces and manipulation into position. This task tests motor behaviour requiring the cooperation of different cognitive functions including motor planning but also mental rotation, working memory and spatial attention to name but a few. It would be intriguing to contrast these results with data collected from tasks requiring hand cooperation in different contexts, to dissociate whether changing the cognitive load can modulate hand as well as action selection.

To conclude, our results provide preliminary evidence to support the role of dorsal fronto-parietal tracts in lateralised hand selection for reaching and grasping movements. While these dorsal white matter structures have already been associated with goal-directed hand movements in monkeys and humans, to our knowledge this study is the first to demonstrate that disrupting their structural asymmetry with unilateral lesions directly alters the choice of hand selected for these movements. This may provide intriguing avenues for future study within the field of motor control and attention, but also for understanding the importance of balance in the relative contributions of each hemisphere toward a single cognitive process.

## Funding bodies

This work was supported by a grant from Associazione Italiana per la Ricerca sul Cancro (AIRC) to L.B and supported by “Regione Lombardia” under the Eloquentstim Project (Por-Fesr, 2014–2020).

